# Enhanced Tumor Control and Hearing Loss Prevention Achieved with Combined Immune Checkpoint Inhibitor and Anti-VEGF Therapy in Vestibular Schwannoma Model

**DOI:** 10.1101/2024.12.29.630658

**Authors:** Simeng Lu, Zhenzhen Yin, Limeng Wu, Yao Sun, Jie Chen, Lai Man Natalie Wu, Janet L. Oblinger, Day Caven Blake, Lukas D. Landegger, Richard Seist, William Ho, Bingyu Xiu, Adam P. Jones, Alona Muzikansky, Konstantina M. Stankovic, Scott R. Plotkin, Long-Sheng Chang, Lei Xu

## Abstract

**Background:** *NF2*-related schwannomatosis (*NF2*-SWN) is a debilitating condition that calls for robust treatment options. The defining feature of *NF2*-SWN is the presence of bilateral vestibular schwannomas (VSs), which grow over time and can result in irreversible sensorineural hearing loss, significantly affecting the quality of life for those affected. At present, there are no FDA-approved medications specifically for treating VS or related hearing loss. VS management involves radiotherapy or surgical resection, while bevacizumab, an anti-vascular endothelial growth factor (VEGF) monoclonal antibody (αVEGF) may be used off-label in *NF2-*SWN to shrink the tumor. However, not all patients respond, and the effect is not always durable. There is a critical need for effective medications that can stop the growth of VS and prevent hearing loss associated with these tumors. While immune checkpoint inhibitors have transformed cancer therapy, their potential has not been thoroughly explored in non-malignant tumors such as VS.

**Methods:** We characterize the effects of anti-PD1 (αPD1) treatment on tumor growth and hearing function in two syngeneic, immune-competent VS models.

**Results:** We demonstrated that combining αVEGF treatment with αPD1 significantly enhances the efficacy of each monotherapy. Specifically, i) αVEGF enhances αPD1 efficacy by normalizing the tumor vasculature to improve drug delivery and immune cell infiltration, and by activating T cell and NK cell anti-tumor cytotoxicity via NKG2D upregulation; and ii) combining αPD1 with αVEGF treatment effectively controls tumors that progressed despite αVEGF treatment.

**Conclusion:** These findings provide a strong foundation for the development of αPD1 with αVEGF combination therapies for patients with *NF2*-SWN.

**Key points:** We filled a critical gap in NF2 research:

1) we characterized the effects of immunotherapy on tumor growth and hearing function in non-malignant vestibular schwannomas

2) We showed combined anti-VEGF and anti-PD1 enhances the efficacy of each monotherapy

**Importance of the study:** Treatment options for patients with *NF2*-SWN are limited or are associated with significant co-morbidities. There are no approved medical treatments for NF2-related tumors. While immune checkpoint inhibitors have transformed cancer therapy, their potential has not been thoroughly explored in non-malignant tumors such as VS. Our work filled this critical gap in *NF2*-SWN research. For the first time, we systemically evaluated ICI efficacy on tumor growth and hearing function in non-malignant schwannomas.

Furthermore, we demonstrated that combining αVEGF treatment with αPD1 significantly enhances the efficacy of each monotherapy. Specifically: i) αVEGF enhances αPD1 efficacy by normalizing the tumor vasculature to improve drug delivery and immune cell infiltration, and by activating T cell and NK cell anti-tumor cytotoxicity via NKG2D upregulation; and ii) combining αPD1 with αVEGF treatment effectively controls tumors that progress despite αVEGF treatment.

Our findings provide a strong foundation for the development of αPD1 with αVEGF combination therapies for patients with *NF2*-SWN.

## Introduction

*NF2*-related Schwannomatosis (*NF2*-SWN) is a dominantly inherited neoplastic syndrome resulting from germline pathogenic variants in the *NF2* tumor suppressor gene. It has an incidence rate of approximately 1 in 61,000 individuals, with nearly complete penetrance at close to 100% ^1,2^. Patients with *NF2*-SWN develop peripheral nerve sheath tumors, characterized by bilateral VSs that arise from cranial nerve VIII. The progressive growth of these VSs can lead to sensorineural hearing loss (SNHL), significant morbidity, and adverse effects on quality of life ^3^. In some cases, large VSs can put pressure on the brainstem, leading to severe complications and even fatality ^4^. Current treatments for progressive VSs include surgery and radiotherapy, both of which carry the risk of further nerve damage and may result in profound deafness, chronic dizziness, and paralysis of the facial and other cranial nerves ^5–7^. There remains a significant unmet medical need for effective treatments that can halt VS growth and prevent SNHL associated with these tumors.

Bevacizumab, a humanized monoclonal antibody that neutralizes VEGF-A, is approved in the UK for the treatment of *NF2*-SWN and has documented benefits in 30-40% of patients, with improvement in hearing or tumor shrinkage ^7–9^. However, several challenging issues remain with bevacizumab therapy: i) some patients do not respond to bevacizumab monotherapy, ii) some patients experience a decrease in tumor growth rate but require additional therapy after long-term treatment, and iii) some patients are unable to tolerate long-term bevacizumab treatment ^10^. Therefore, there is a need for combination regimens with bevacizumab or novel alternative therapies to control VS tumors that progress despite bevacizumab treatment.

Immune checkpoint inhibitors (ICIs), including monoclonal antibodies that target programmed cell death protein-1 (PD-1), PD ligand 1 (PD-L1), and cytotoxic T-lymphocyte-associated protein 4 (CTLA-4), work by blocking immune inhibitory signals to enhance T cell activation and functionality. This process leads to improved antitumoral immunity and better prognostic outcomes ^11,12^. ICIs have received FDA-approval for a variety of solid tumors, either as standalone treatments or in conjunction with therapies such as chemotherapy^13^. However, the advantages of ICIs are currently confined to a specific group of patients, and their therapeutic potential in non-malignant schwannomas has yet to be investigated.

VS tumors contain a significant population of T cells ^14^. However, many of these infiltrating CD4^+^ and CD8^+^ T cells express PD-1 ^15^ and display a transcriptome signature indicative of CD8^+^ T cell senescence ^16^. Additionally, PD-1 and its ligand B7-H1 are expressed in VS tumors ^14,17^. Despite these findings, there has been limited research on the therapeutic potential of ICIs in VS. Notably, an anti-PD1 (αPD1) antibody was shown to modestly slow schwannoma growth in a subcutaneous mouse model ^18^, and αPD1 salvage therapy resulted in tumor growth inhibition in a patient with recurrent VS ^19^. These insights indicate that ICIs could represent a promising treatment approach for VS.

In this study, we aim to address two questions: 1) can anti-VEGF (αVEGF)-induced vessel normalization enhance intratumoral delivery of ICI drug and immune effector cells, and thus augment the ICI efficacy in VS? and 2) can αPD1 serve as an effective alternative for patients unresponsive to or unable to tolerate bevacizumab?

## Materials and Methods

### Cell lines

Mouse *Nf2*^-/-^ Schwann cells were maintained in 10% fetal bovine serum (FBS)-containing Schwann cell medium, which includes Schwann cell growth supplement (SCGS, ScienCell)^20^. Mouse SC4 Schwannoma cells (gift from Dr. Vijaya Ramesh, Massachusetts General Hospital) were maintained in 10% FBS-containing Dulbecco’s Modified Eagle Medium (DMEM, Corning)^21,22^.

### Animal models

All animal procedures were conducted in accordance with the Public Health Service Policy on Humane Care of Laboratory Animals and received approval from the Institutional Animal Care and Use Committee of the MGH. *Nf2^-/-^*and SC4 tumors were inoculated into immune-competent C57/FVB mice. In all animal experiments, we utilized male and female mice aged 8-12 week-old mice to ensure sufficient statistical power and to examine any potential sex-related differences. Both sexes were represented equally (1:1 ratio), and we matched mice by age and sex for each experimental condition. Given that schwannomas develop in the vestibular nerve and peripheral nerves in patients, we employed two mouse models by injecting tumor cells into the cerebellopontine angle (CPA) and into the sciatic nerve of the mice.

#### Cerebellopontine angle (CPA) model

To simulate the intracranial microenvironment of VSs, tumor cells were injected into the CPA region of the right hemisphere ^23,24^. Each mouse received an implantation of 1 μl of tumor cell suspension containing 2,500 cells.

#### Sciatic nerve schwannoma model

To mimic the microenvironment associated with peripheral schwannomas, we injected tumor cells into the mouse sciatic nerve of the mice ^21^. We used a Hamilton syringe to slowly inject 3 μl of tumor cell suspension (5×10^4^ cells) over the course of 45-60 seconds, placing the inoculum under the sciatic nerve sheath to avoid any leakage.

### Treatment protocols

In the sciatic nerve model, treatment begins once the tumor reaches a diameter of 3 mm. In the CPA model, treatment starts when the blood concentration of the *Gaussia* luciferase reporter gene (Gluc) reaches 1×10^4^ RLU (relative luminescence unit). For αPD1 treatment, the αPD1 antibody or isotype control IgG (200 μg/mice, BioXCell) was administrated *i.p.* every 3 days for a total of 4 dosages. For αVEGF treatment, the αVEGF antibody B20.4.4 (B20, 2.5 mg/kg, Genentech), which neutralizes both human and mouse VEGF, was administered *i.p.* on a weekly basis ^21^.

### Measurement of tumor growth

To track tumor growth in the CPA model, both tumor cell lines were transfected with lentivirus carrying the secreted Gluc reporter gene, and plasma Gluc was measured as previously described ^24–27^. In brief, 13 μl of whole blood was collected from the tail vein and immediately mixed with 5 μl 50mM EDTA to prevent clotting. The blood sample was then transferred to a 96-well plate, where Gluc activity was assessed using a plate luminometer (GloMax 96 Microplate Luminometer, Promega). The luminometer was configured to automatically inject 100 µl of 100 mM coelenterazine (CTZ, Nanolight) in PBS, with photon counts recorded for 10 sec. The size of the tumor in the sciatic nerve was measured with calipers every 3 days until the tumors reached a diameter of 1 cm.

### Audiometric testing in animals

Auditory brainstem responses (ABRs) were assessed as previously outlined^24^. In brief, the animals were anesthetized with an intraperitoneal injection of ketamine (0.1 mg/g) and xylazine (0.02 mg/g). The tympanic membrane and middle ear were examined microscopically for any signs of otitis media, and all animals showed well-aerated middle ears. ABRs were collected using subdermal needle electrodes positioned as follows: the positive electrode was placed on the inferior aspect of the ipsilateral pinna, the negative electrode at the vertex, and the ground electrode at the base of the tail. The responses were amplified by a factor of 10,000, filtered within a range of 0.3–3.0 kHz, and averaged over 512 repetitions for frequency and sound level that were used in the DPOAE tests. Data acquisition was performed using custom LabVIEW software running on a PXI chassis from National Instruments Corp. For each frequency tested, the auditory threshold was determined as the lowest stimulus level at which consistent peaks could be visually identified. If no auditory threshold was observed, a value of 85 dB was recorded, representing 5 dB above the highest level tested.

### Analysis of tumor-infiltrating immune cells

To analyze the immune cells that infiltrate the tumor, tumor lysates were treated with PBS and stained for flow cytometry using several antibodies: anti-CD45 (clone 30-F11), anti-CD4 (clone RM4-5), anti-CD8 (clone 53-6.7), anti-NK1.1 (clone PK136), anti-Gr1 for MDSC (clone RB6-8C5), and anti-CD11b (clone m1/70). Please refer to Supplementary Table 1 for detailed information about the antibodies used.

### Gene expression analysis

#### Single-cell RNA-Sequencing (scRNA-seq)

scRNA-seq was conducted on human VSs obtained from *NF2*-SWN patients who provided informed consent, including those who received bevacizumab treatment, in accordance with the Human Subjects protocol approved by the Ohio State University Institutional Review Board-approved Human Subjects protocol for scRNA-seq. Briefly, fresh surgically removed VS tissues were placed in DMEM or saline and transported to the lab for preparing single-cell suspension as previously described (*28a*).

Dissociated cells were cryopreserved in 10% dimethyl sulfoxide in DMEM containing 10% FBS until processing on the 10X Genomics’ Chromium Single Cell platform and sequencing on a NovaSeq 6000 System (Illumina) at a depth of approximately 400 million reads per sample. For computational analysis, we used Cellranger v5.0.1 to align reads to the hg19 human reference sequence. For each sample dataset, unsupervised clustering was performed using R package Seurat (version 4)(https://www.satijalab.org/seurat, https://www.github.com/satijalab/seurat)^28^. Following normalization using Seurat’s NormalizeData function, highly variable genes were identified and used for principal components analysis. Then, we performed clustering using graph-based clustering and visualized using Uniform Manifold Approximation and Projection (UMAP) with Seurat function RunUMAP. To identify the cell type in tumor sample datasets, we input marker gene lists generated by the Seurat FindMarkers function in Toppgene to identify the top cell type makers or cell identities. Differentially expressed genes (DEGs) in a given cell type compared with all other cell types were determined with the FindAllMarkers function in the Seurat package.

#### Quantitative RT-PCR

Changes in the mRNA levels were quantified with qPCR, using SYBR Green-based protocol ^29^. All qPCR and analysis were performed on a Stratagene MX 3000 qPCR System with the operating MXPro qPCR software (Stratagene)^30^.

#### Western Blot

Thirty micrograms of protein per sample were separated on 10% SDS-polyacrylamide gels ^31^. Membranes were blotted with antibodies against perforin (1:500) and granzyme B (1:500) (Cell Signaling Technology). Membranes were blotted with the antibody against β-actin for an equal loading control (1:5000, Sigma)^32^.

#### ELISA

Plasma or protein obtained from snap-frozen tumors was diluted to a concentration of 2 μg/μl based on protein assays. Levels of inflammatory cytokines in the mice were measured using mouse multiplex enzyme-linked immunosorbent assay plates, in accordance with the manufacturer’s guidelines (Meso-Scale Discovery). Each sample was tested in triplicate ^29,33^.

### *In vitro* cytotoxicity assay

CD8^+^ T cells were isolated from treated tumors, and polyclonal NK cells were isolated from spleens using the T cell and NK Cell Isolation Kits and MACS columns (Miltenyi Biotech) following the manufacturer’s protocol ^34^. Experiments were done when the purity of CD8^+^ T cells and NK cells was above 90% as determined by flow cytometry. Cultured mouse lymphoma YAC-1 cells or schwannoma tumor cells were used as the target cells and labeled with calcein AM (10 nM for 1×10^6^ cells/ml, Becton Dickson). After overnight incubation, 1×10^4^ cells/well were seeded in a U-bottom 96-well plate, 100 μl/well, protected from light. The effector cells were added in triplicate at the effector:target (E:T) cell ratio of 10:1.

Target cells only were plated for baseline fluorescent reading, and target cells in lysis buffer (1% NP40) were included as maximum cell lysis. After incubation for 6 hours, the supernatants were collected, and fluorescent intensity was measured using a fluorescent plate reader. Cytolysis was assessed by measuring the release of calcein AM from target cells as described previously ^35^: effector cell cytotoxicity (%) = 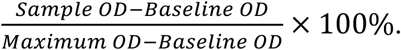.

### Histological staining

To evaluate tumor cell proliferation and apoptosis, sections of tumor tissues were stained with proliferating cell nuclear antigen (PCNA, 1:1000, Abcam) and TUNEL (ApopTag®, EMD Millipore).

Microvessel density (MVD, CD31, 1:200, Millipore) and myofibroblast infiltration (α-Smooth Muscle - Cy3™, αSMA-Cy3, 1:200, Sigma-Aldrich) were determined in frozen sections (8 μm). To evaluate the percentage of perfused vessels, 50 μl of FITC-lectin (2 mg/kg) were injected *i.v.* 10 minutes prior to sacrifice to identify perfused vessels. Ten minutes later, tumors were harvested and embedded in frozen sections (20 μm).

Endothelial cell labeling by FITC-lectin demonstrates perfused blood vessels, and CD31^+^ staining demonstrates all blood vessels in solid tumors; the percentage of FITC-lectin^+^/CD31^+^ vessels was quantified using Image J software. To fluorescently label αPD1 antibodies, Alexa Fluor™ 647 Protein Labeling Kit (A20173, ThermoFisher Scientific) was used following the manufacturer’s protocol. The labeled αPD1 was processed using the purification columns provided in the kit to remove the unbound dye and purify the labeled antibody. Four specimens of normal peripheral nerve were obtained postmortem and used as controls. Appropriate positive and negative controls were used for all stains ^33^. Histological analysis using digital quantitative image analysis was performed using the open-source software ImageJ. Positive staining in 20 random fields/slides was quantified via automated built-in functions based on fluorescent pixel intensity after establishing a threshold to exclude background staining.

### Statistical analyses

To assess whether growth curves significantly differed from one another, we log-transformed the data, fitted linear regression models to each growth curve, and compared the slopes of the regression lines using an F-test from ANCOVA. Survival analyses and curve generated were performed using the KM method. Differences in tumor growth between the two groups were evaluated using either the two-tailed Student’s t-test or the two-tailed Mann-Whitney *U* test. For comparing gene expression, we utilized mixed-effect models to account for the correlation structure. P values underwent multiple-test correction through the Benjamini-Hochberg False Discovery Rate (FDR) method. The Benjamini-Hochberg adjustment for multiple comparisons was implemented. All calculations were conducted using GraphPad Prism Software version 6.0 and Microsoft Excel 2010.

## Results

### αVEGF treatment normalizes tumor vasculature, and increases αPD1 drug delivery and intratumoral infiltration of immune effector cells in the mouse schwannoma model

To address our first question of whether αVEGF-induced vessel normalization enhances intratumoral delivery of ICI drug and immune effector cells and thus augment the ICI efficacy in VS, we evaluated how αVEGF treatment affects tumor vasculature in the *Nf2^-/-^* CPA model. αVEGF treatment did not significantly reduce microvessel density (MVD, Figure 1A-B), but significantly increased the fraction of pericyte-covered vessels, indicating these tumor blood vessels are structurally similar to normal vessels (Figure 1C). Next, we measured the fraction of perfused vessels to determine whether the structural normalization of tumor vessels leads to normalized vessel perfusion. We injected FITC-lectin *i.v.* to identify perfused tumor vessels and stained for CD31 to detect the total number of blood vessels (Figure 1D). In concert with the blood vessel structural normalization, αVEGF treatment increased the percentage of perfused vessels (Figure 1E). These data indicate that αVEGF treatment normalized schwannoma vasculature.

**Figure 1.**
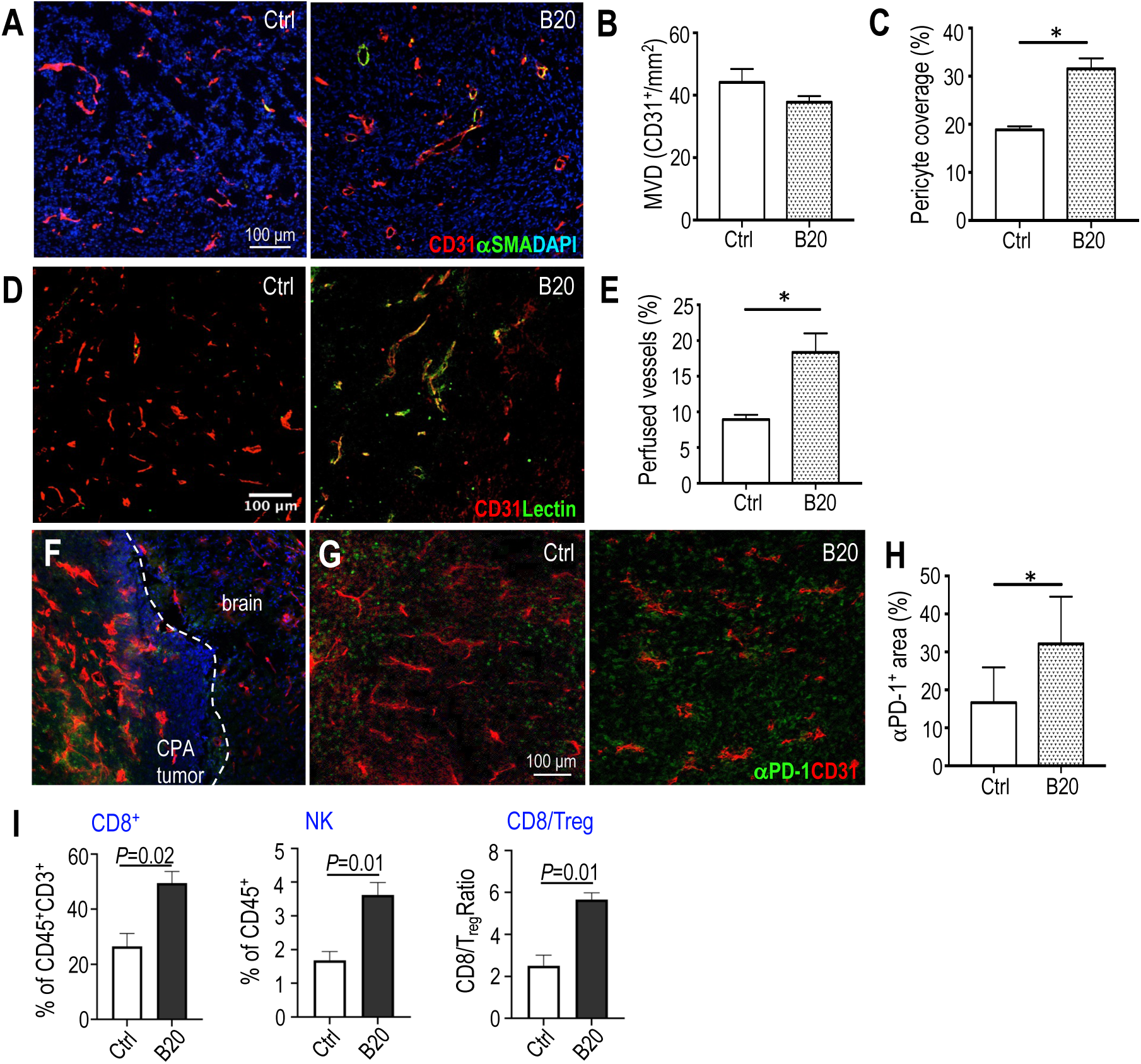
αVEGF treatment normalizes tumor vasculature, increases αPD1 drug delivery and immune effector cell intratumoral infiltration in the mouse schwannoma model. **(A)** Representative immunofluorescent (IF) staining image of tumor blood vessel endothelial cells (CD31^+^, red), pericyte (αSMA^+^, green) in *Nf2^-/-^* tumor (DAPI, blue). **(B)** Microvessel density (MVD) was quantified by manual counting of CD31^+^ (red) vessels/mm^2^. **(C)** The percentage of pericyte covered blood vessels (αSMA^+^CD31^+^, yellow) of total blood vessels (CD31^+^, red) was quantified using ImageJ software. **(D)** Representative IF staining images of perfused vessels (FITC-lectin^+^/green, yellow), and total blood vessels (CD31^+^, red). **(E)** The percentage of perfused vessels was quantified using ImageJ software. Representative images of the distribution of FITC-labeled αPD1 antibody (green) in: **(F)** the brain and CPA tumors, **(G)** control-IgG and B20-treated tumors. Red, CD31^+^ tumor blood vessels; blue, DAPI. **(H)** Quantification of the fraction of tumor area positive for αPD1 antibody was performed using ImageJ. **(I)** Flow cytometry analysis of CD8^+^ T cell, NK cells and CD8/T_reg_ ratio in *Nf2^-/-^* tumors. For histological staining, samples from n=4 mice/group were used, 20 random areas were imaged and quantified. Image quantification is presented as mean±SD and analyzed using Student’s t-test and the Mann-Whitney test. *P<0.01.

Next, to explore whether αVEGF-improved vessel perfusion can increase the delivery of large molecular weight antibodies, we labeled the αPD1 antibody with a green fluorescein tag ^36^ and injected it *i.p.* into control IgG-or αVEGF-treated mice in the *Nf2^-/-^* CPA model. Tumor-bearing brains were collected 24 hours later. We observed significant angiogenesis at the peritumoral margin, and the green fluorescent αPD1 antibody predominantly accumulated within the tumor, with minimal presence in the surrounding brain tissue (Figure 1F). In αVEGF-treated tumors, we observed a significantly higher intratumoral fluorescence signal than in control IgG-treated tumors (Figure 1G), indicating increased delivery of αPD1 antibody (Figure 1H).

Then, using flow cytometry to analyze the intratumoral immune cell population, we found that αVEGF treatment significantly increased the number of tumor-infiltrating cytotoxic CD8^+^ T cells and NK cells, as well as increased the ratio of CD8^+^ T cells over immune suppressive T_reg_ (CD8^+^/T_reg_, Figure 1I).

### Combined αVEGF treatment enhances αPD1 efficacy in the mouse schwannoma models

The αVEGF-induced increased delivery of αPD1 antibody and immune effector cells provides the rationale to test if αVEGF treatment could enhance αPD1 efficacy. We treated mice bearing *Nf2^-/-^* tumors in both sciatic nerve and CPA models with: i) control IgG, ii) αPD1, iii) αVEGF, or iv) αPD1+αVEGF RT (**Figure 2A**). In the sciatic nerve model, combined αVEGF treatment enhanced the anti-tumor effect of αPD1 and significantly delayed tumor growth compared to αPD1 and αVEGF monotherapies (**Figure 2B**). This enhanced efficacy from the combination treatment was also observed in a second schwannoma, SC4 model (Supplemental Figure 1). In the *Nf2^-/-^*CPA model, combined αVEGF treatment significantly prolonged animal survival – with 76% of mice surviving ≥100 days, compared to 32% of long-survivor mice in the αPD1 monotherapy group (**Figure 2C**). In the CPA tumors collected from mice in the combination group, we observed more apoptotic tumor cells (TUNEL^+^) and fewer proliferating tumor cells (PCNA^+^) compared to those in the control or monotherapy groups (Supplemental Figure 2A-B).

**Figure 2.**
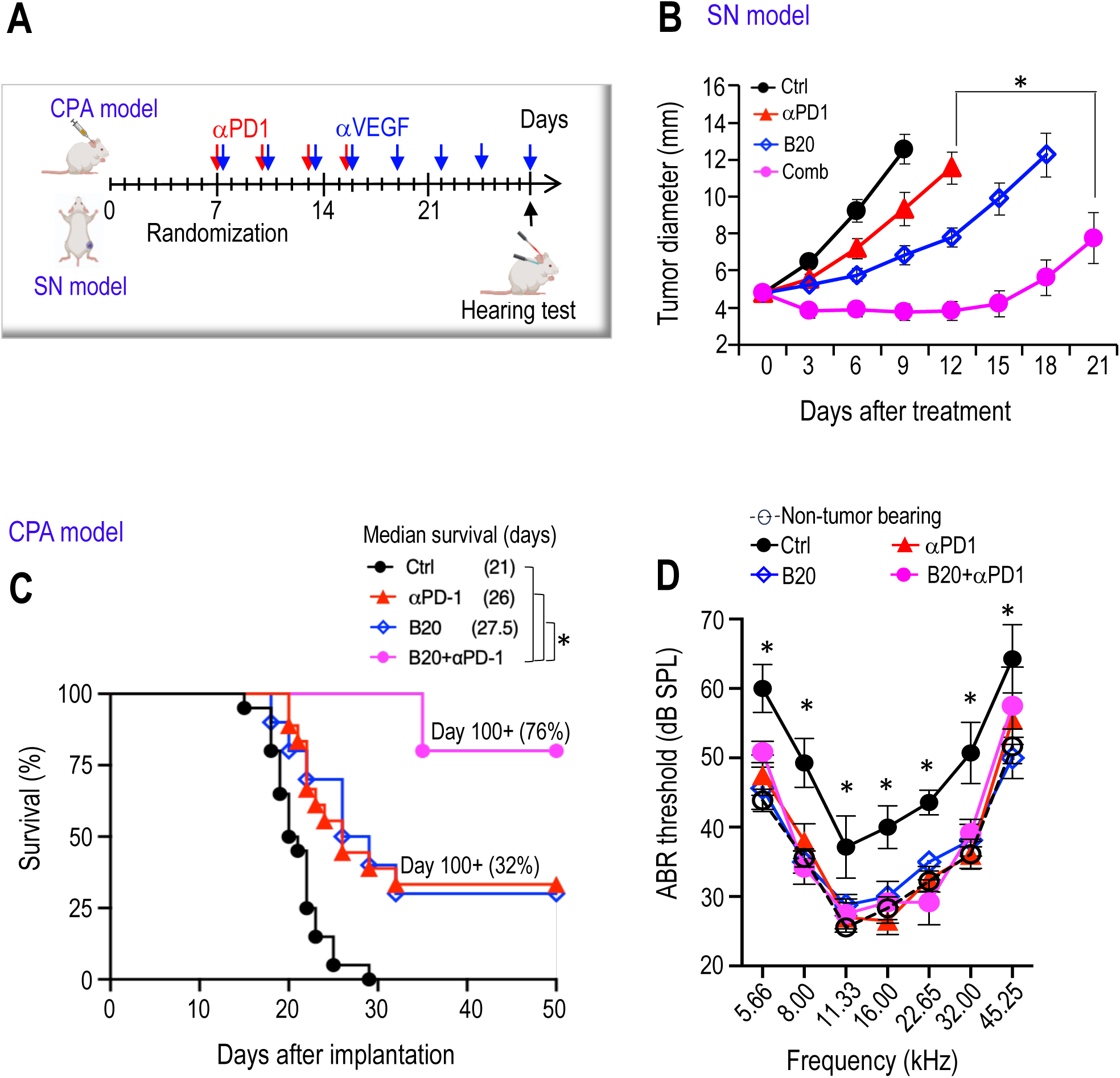
Combined αVEGF treatment enhances αPD1 efficacy in the mouse schwannoma models. **(A)** Schematic and timeline of combined αVEGF and αPD1 treatment in the CPA and SN mouse schwannoma models. **(B)** Tumor diameter measured by caliper in the SN model. **(C)** Kaplan-Meir survival curve of mice bearing *Nf2^-/-^* tumor in the CPA model. **(D)** ABR threshold of mice bearing *Nf2^-/-^* tumor in the CPA model. All animal studies are presented as mean±SEM, N=6 mice/group, and representative of at least three independent experiments. In vivo study significance was analyzed using a Student’s t-test. *P<0.05.

The effects of immunotherapy on hearing function remain to be systemically evaluated. We first evaluated potential ototoxicity from αPD1 treatment in non-tumor bearing mice by measuring auditory brainstem evoked response (ABR), which represents the summed activity of the auditory nerve and central auditory nuclei. In non-tumor-bearing mice, three doses of αPD1 treatment did not change the ABR threshold in mice 21 days post-treatment (Supplemental Figure 3A-B), demonstrating that αPD1 does not cause acute ototoxicity. In mice bearing *Nf2^-/-^* tumors in the CPA region, αVEGF prevented tumor-induced hearing loss (blue line), reproducing the hearing benefit from bevacizumab treatment observed in patients with *NF2*-SWN. αPD1 monotherapies (red line), as well as their combination with αVEGF treatment (pink line), restored the ABR threshold to normal levels observed in non-tumor-bearing mice. No treatment demonstrated superiority over the others (Figure 2D).

### αVEGF-enhanced αPD1 efficacy is mediated by CD8^+^ T cells and NK cells

Using flow cytometry to analyze the intratumoral immune cell population, we found that both αPD1 and αVEGF monotherapies significantly increased the number of tumor-infiltrating cytotoxic CD8^+^ T cells and NK cells compared to the control group. Additionally, αPD1 monotherapy reduced the number of immune-suppressive myeloid-derived suppressor cells (MDSCs). Compared to the αPD1 monotherapy, combined αVEGF treatment further increased the infiltration of effector CD8^+^ T cells and NK cells, while reducing MDSCs (Figure 3A). In our analysis of the tumor secretome, we observed that αPD1 treatment elevated levels of IFN-γ and TNF-α, cytokines known to synergistically activate T cells and produced by activated T cells. Furthermore, combined αVEGF and αPD1 treatment significantly increased IFN-γ and TNF-α production compared to αPD1 monotherapy (Figure 3B).

**Figure 3.**
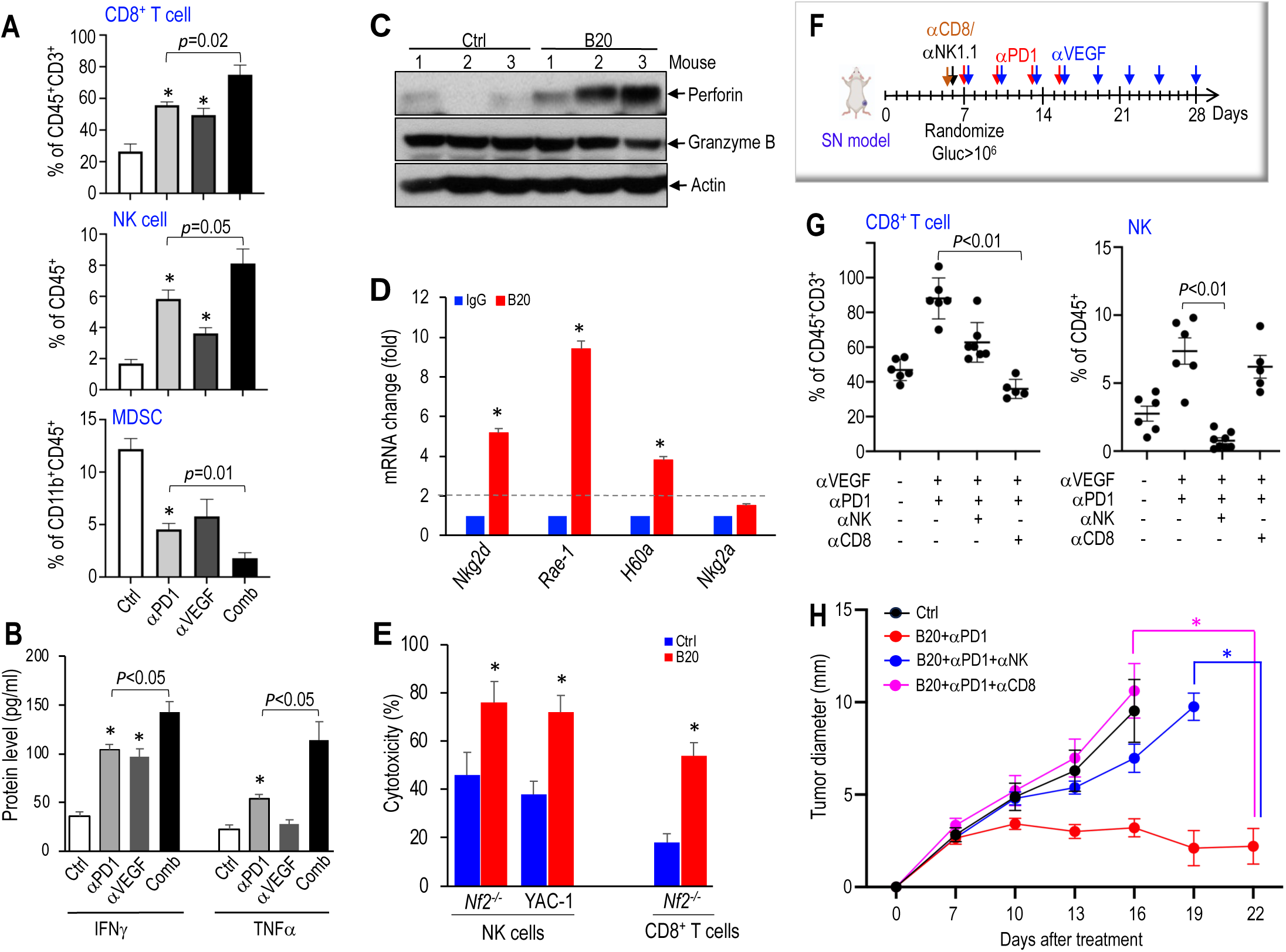
αVEGF-enhanced αPD1 efficacy is mediated by CD8^+^ T cells and NK cells. **(A)** Flow cytometry analysis of the number of CD8^+^ T cell, NK cell and MDSC in *Nf2^-/-^* tumor from different treatment groups (N=3 tumors/group). **(B)** ELISA of murine IFNγ and TNFα in in *Nf2^-/-^* tumor from different treatment groups (N=3 tumors/group). **(C)** Western blot of tumor tissues. **(D)** qRT-PCR analysis of murine *Nkg2d, Nkg2a, Rae-1, and H60a* mRNA in *Nf2^-/-^* tumor (N=3 tumors/group). **(E)** *In vitro* T cell and NK cell cytotoxicity assay. **(F)** Schematic and timeline of CD8 T cell and NK cell depletion in *Nf2^-/-^* sciatic nerve model. **(G)** Flow cytometry analysis of the number of CD8^+^ T cell and NK cell in *Nf2^-/-^* tumors from different treatment groups. **(H)** Tumor diameter measured by caliper in the SN model. Flow cytometry, ELISA, qPCR and cytotoxicity studies are presented as mean ± SD, and analyzed using Student’s t-test and the Mann-Whitney U test. All animal studies are presented as mean±SEM, N=8 mice/group, and representative of at least three independent experiments. *In vivo* study significance was analyzed using a Student’s t-test. *P<0.01.

These findings suggest that in addition to bringing in more CD8^+^ T cells and NK cells into the tumor, αVEGF treatment also played a role in activating these immune effector cells.

We next investigated how αVEGF affects the activation of CD8^+^ T cells and NK cells. Perforin and granzyme B are critical in both CD8^+^ T cell and NK cell cytotoxicity ^37,38^. We observed that αVEGF treatment significantly induced perforin expression but did not change the granzyme B level in *Nf2^-/-^* tumors (Figure 3C). In *Nf2^-/-^* tumors, αVEGF treatment significantly increased the expression of natural killer group 2D (*Nkg2d)*(Figure 3D). NKG2D is an activating receptor expressed on NK cells, CD8^+^ T cells, and activated CD4^+^ T cells, which promotes NK cell cytotoxicity ^39^ and augments T cell receptor-mediated activation of CD8^+^ T cells ^40^. In mice, NKG2D binds to ligands, including RAE-1 and H60A ^41,42^. αVEGF treatment also significantly increased the expression of NKG2D ligands *Rae-1* and *H60a*, but did not affect the expression of the inhibitory receptor *Nkg2a* (Figure 3D).

To investigate whether αVEGF treatment regulates the cytotoxic function of NK cells and CD8^+^ T cells, we treated *Nf2^-/-^* tumor-bearing mice with control IgG or B20 for 21 days. Following sacrifice, we isolated: i) tumor-associated CD8^+^ T cells from control- and B20-treated mice, and ii) spleen NK cells, given the low number of tumor-infiltrating NK cells. CD8^+^ T cells were stimulated with IL-2 (50 IU/ml) and co-cultured with calcein AM-labeled *Nf2^-/-^*tumor cells at an effector-to-target ratio (E:T) of 10:1. Similarly, NK cells were co-cultured with calcein AM-labeled *Nf2^-/-^* tumor cells and YAC-1 cells, which are sensitive to NK cell-mediated cytotoxicity, at the same E:T ratio. After 8 hours of co-culture, target cell lysis was quantified by measuring the fluorescent intensity in the supernatant. CD8^+^ T cells and NK cells isolated from αVEGF-treated mice exhibited significantly enhanced cytotoxicity against both *Nf2^-/-^* tumor cells and YAC-1 target cells compared to effector cells isolated from control IgG-treated mice (Figure 3E).

These data suggest that αVEGF-induced influx and activation of immune effector CD8^+^ T cells and NK cells are key immunologic mechanisms mediating the enhanced efficacy of combined αVEGF and αPD1 treatment. To test this hypothesis, we depleted mice of CD8^+^ T cells or NK cells before treating them with the combination therapy (Figure 3F). The treatment of mice with anti-CD8 depletes CD8^+^ T cells, and the treatment of anti-NK1.1 specifically reduced the number of NK cells as confirmed by flow cytometry (Figure 3G).

Depletion of CD8^+^ T cells completely abolished the tumor control benefit of the combination therapy, while depletion of NK cells significantly reduced the therapeutic benefit (Figure 3H).

### scRNA-Seq reveals elevated cytotoxic profiles of CD8^+^ T and NK cells in VS from an *NF2*-SWN patient treated with bevacizumab

To profile the T cells and NK cells in VSs and characterize their changes with or without bevacizumab treatment, we dissociated fresh VS tissues from *NF2*-SWN patients who were either bevacizumab-naïve (n=3) or treated with bevacizumab for nearly 10 years (n=1). Single-cell transcriptomic profiling was performed using the 10X Genomics platform. Within the VS tissues, cell clusters were partitioned into major cell types, including Schwann cell (SC) lineage, macrophages, lymphocytes, fibroblasts, and other stromal cells (Chang LS, et al., manuscript in preparation). We further subclustered the lymphocyte compartment and identified distinct lymphocyte-derived cell states, including CD8^+^ cytotoxic T cells, early naïve T cells, CD8^+^ PD-1^+^ T cells, regulatory T cells, and NK cells in naïve VS (Figure 4A).

**Figure 4.**
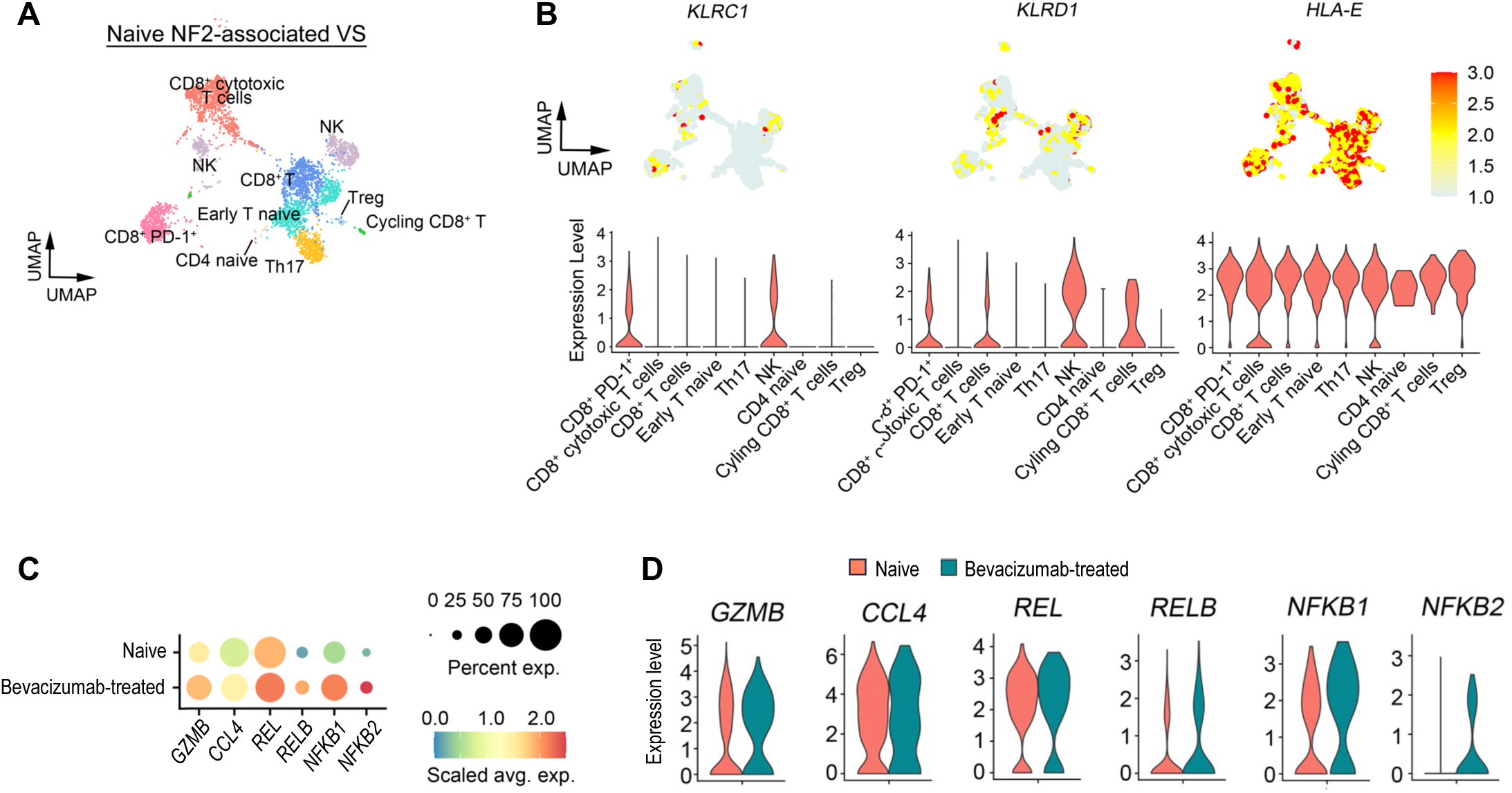
scRNA-Seq reveals elevated cytotoxic profiles of CD8^+^ T and NK cells in VS from an NF2 patient treated with bevacizumab. **(A)** UMAP visualization of various lymphocyte subpopulations from treatment-naïve NF2-associated VS. Colors represent assigned cell types. **(B)** The gene expression (top) and violin plots (bottom) of *KLRC1, KLRD1* and *HLA-E* expression within lymphocyte subpopulations in naïve *NF2*-associated VS. **(C)** The dot plot of cytotoxicity marker gene expression in NK cells in the VSs with or without bevacizumab treatment. The color scale represents scaled average expression of the indicated marker genes in each tumor type, and the size of the circle indicates the proportion of cells expressing each marker gene. **(D)** The violin plots of cytotoxicity marker gene expression in NK cells in the VS with or without bevacizumab treatment.

In NK cells and CD8^+^ T cells from naïve VSs, we observed elevated RNA expression of i) *KLRC1*, which encodes NKG2A, a major inhibitory receptor expressed on NK cells and CD8^+^ T cells ^43,44^; ii) *KLRD1*, which encodes CD94, forming a heterodimer with NKG2A; and iii) *HLA-E*, the ligand for NKG2A:CD94 heterodimer, which sends a strong inhibitory signal regulating NK cell and CD8^+^ T cell cytotoxic activity (Figure 4B)^45,46^. These scRNASeq findings indicate that in the VS tumor microenvironment, CD8^+^ T cells and NK cells are in a suppressed functional state.

In VS treated with bevacizumab, compared to naïve VS, we observed increased expression of CD8^+^ T cell and NK cell cytotoxicity markers, including i) Granzyme B (*GZMB)*, the key mediator of NK cell and CD8^+^ T cell cytotoxicity ^47^, ii) *CCL4*, chemokine recruiting NK cells and T cells ^48^, iii) NFκB pathway genes*, including REL, NFκB1,* and *NFκB2*, which play a crucial role in regulating the cytotoxic activity of both NK cells and T cells ^49^(Figures 4C-D). These scRNASeq findings indicate that bevacizumab treatment enhanced the cytotoxic function of CD8^+^ T and NK cells.

### αPD1 can control the growth of tumors that progress despite αVEGF treatment in Schwannoma models

We next performed an experiment to address our second question: can αPD1 serve as an effective alternative for patients who are unresponsive to or unable to tolerate bevacizumab? We implanted mouse *Nf2^-/-^* cells in both the sciatic nerve and the CPA models and treated the mice with 3 doses of αVEGF. Then, the mice were randomized into three groups, receiving: i) continued αVEGF monotherapy, ii) combined αPD1 with αVEGF treatments, or iii) switched to αPD1 monotherapy, with αVEGF discontinued (Figure 5A).

**Figure 5.**
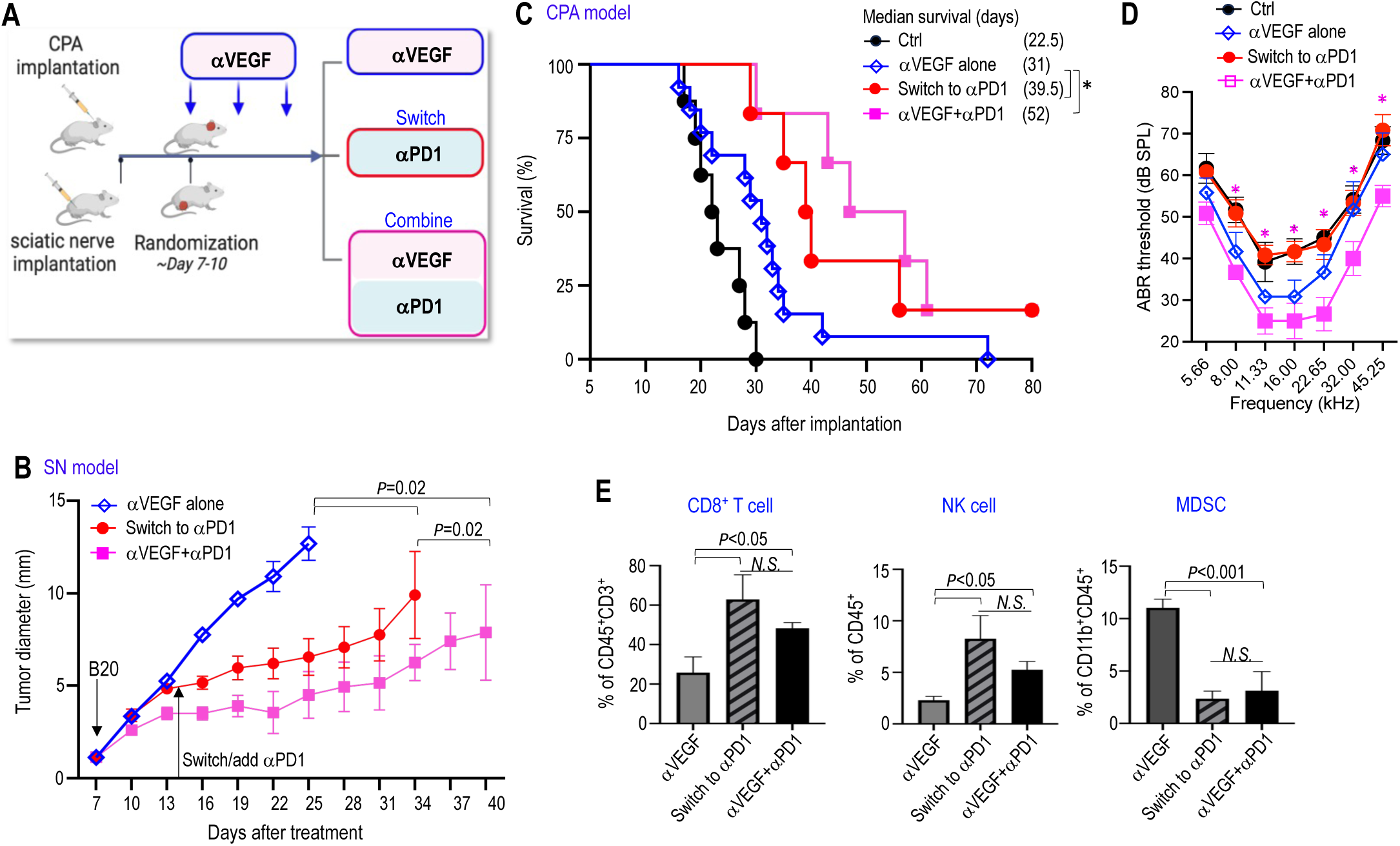
αPD1 can control the growth of tumors that progress despite αVEGF treatment in mouse schwannoma models. **(A)** Schematic and timeline of treatment in the *Nf2^-/-^* CPA models. **(B)** Tumor diameter measured by caliper in the SN model. **(C)** Kaplan-Meir survival curve of mice bearing *Nf2^-/-^* tumor in the CPA model. **(D)** ABR threshold of mice bearing *Nf2^-/-^* tumor in the CPA model. **(E)** Flow cytometry analysis of the number of CD8^+^ T cell, NK cell and MDSC in *Nf2^-/-^* tumor from different treatment groups (N=3 tumors/group). Flow cytometry studies are presented as mean ± SD, and analyzed using Student’s t-test and the Mann-Whitney U test. All animal studies are presented as mean±SEM, N=8 mice/group for tumor growth and survival studies, N=6 mice/group for hearing test, and are representative of at least three independent experiments. In vivo study significance was analyzed using a Student’s t-test. *P<0.01.

In the sciatic nerve model, both switching to αPD1 monotherapy and combining αPD1 with αVEGF significantly delayed tumor growth compared to αVEGF alone, with the combination therapy demonstrating the greatest efficacy (Figure 5B). In the CPA model, while discontinuing αVEGF and switching to αPD1 (red line) only modestly prolonged animal survival, it completely lost the hearing benefit from αVEGF treatment. More importantly, the combination of αPD1 with αVEGF led to the most significant survival benefit, and further prevented tumor-induced hearing loss compared to αVEGF alone (Figure 5C-D). Flow cytometry analysis revealed that both switching to αPD1 and combining αPD1 with αVEGF significantly increased intratumoral CD8^+^ T cells and NK cells while reducing immune suppressive MDSCs compared to αVEGF monotherapy (Figure 5E). These findings suggest that αPD1 can serve as an effective alternative or combination therapy to αVEGF in the treatment of VS. Furthermore, combination therapy provide superior efficacy in both tumor control and hearing preservation.

## Discussion

Our long-term goal is to develop an effective therapy for NF2 patients that not only controls VS tumor growth but also preserves hearing. Using schwannoma mouse models, we have achieved several key findings: i) we conducted the first comprehensive investigation of the immune checkpoint inhibitor αPD1 as a potential treatment for non-malignant VS, characterizing its effects on tumor growth; (ii) we evaluated αPD1’s effect on hearing preservation, which had not been previously explored. More importantly, we report here that combining αVEGF treatment with αPD1 significantly enhances the efficacy of each monotherapy. Specifically, iii) αVEGF treatment, via improving drug delivery and facilitating intratumoral infiltration of immune effector cells, enhanced αPD1 efficacy; and iv) combining αPD1 with αVEGF treatment effectively controlled tumors that progressed despite αVEGF treatment. These findings provide a strong foundation for the development of αPD1 with αVEGF combination therapies for patients with NF2.

ICIs have revolutionized cancer treatment; however, their application to non-malignant schwannomas and their therapeutic potential on hearing preservation remains unexplored. Our study filled this gap, making several discoveries: First, we characterized the effects of αPD1 on VS tumor growth, marking the first comprehensive investigation of an ICI as a potential treatment for non-malignant VS. A previous study using a subcutaneous mouse model reported that αPD1 antibody modestly delayed schwannoma growth ^18^. As a step further, we used two anatomically correct models of schwannoma: i) the CPA model, which reproduces the intracranial microenvironment and recapitulates tumor-induced hearing loss, and ii) the sciatic nerve schwannoma model, which reproduces the nerve microenvironment of peripheral nerve schwannomas. Our results demonstrate that immune checkpoint molecules are valid targets with αPD1 treatment inhibiting schwannoma growth. In the clinic, a case report of αPD1 salvage therapy showed tumor growth arrest in one patient with recurrent VS ^19^.

Our preclinical findings, together with this case report, provide the rationale for future translational studies to better characterize the efficacy of ICIs in patients with VS. Second, we evaluated the effects of αPD1 on hearing preservation, which has not been previously investigated. We first investigated potential ototoxicity from αPD1 treatment. Debilitating and sometimes life-threatening immune-related adverse events (irAEs) can result from aberrant activation of T cell responses following ICIs therapy ^50^. A growing body of research indicates that inflammation plays a critical role in hearing loss ^51,52^. In our mouse model, we observed no acute ototoxic effects from αPD1 treatment and no change in hearing function for up to 3 weeks post-treatment. These findings suggest that αPD-1 may have a manageable safety profile in terms of ototoxicity; however, close monitoring of inflammatory biomarkers in future clinical studies will be necessary to assess the safety of ICI therapy in VS treatment.

The benefits of ICIs are limited to a subset of patients, with efficacy reported in fewer than 20-30% of patients with non-small cell lung cancer, renal cell carcinoma, and melanoma ^53^. In these cancers, the density and spatial distribution of immune infiltrates in tumors are significantly associated with patient survival and response to immune therapy ^54–56^. Abnormal vascular perfusion impedes the intratumoral infiltration of immune cells ^57,58^. In NF2 mouse models, we previously showed that αVEGF normalizes schwannoma vasculature and improves vessel perfusion ^21^. This prompted us to investigate whether αVEGF-improved vessel perfusion could enhance ICI drug delivery and immune cell infiltration, thereby augmenting ICI efficacy in VS? In schwannoma models, we found that αVEGF enhances αPD1 efficacy by i) normalizing the tumor vasculature to improve drug delivery and immune cell infiltration, and ii) by activating T cell and NK cell anti-tumor cytotoxicity via NKG2D upregulation. As a result, the combination treatment demonstrates greater efficacy.

Bevacizumab, a humanized monoclonal antibody that neutralizes VEGF-A, is approved in the UK for the treatment of *NF2*-SWN and has documented benefits in 30-40% of NF2 patients, with improvement in hearing or tumor shrinkage ^9^. However, significant challenges remain: i) not all patients respond to bevacizumab, ii) hearing preservation is often not durable in responders, and iii) some patients may be unable to tolerate long-term treatment due to adverse effects ^10^. Therefore, there is a need for combination regimens or novel therapies to control VS tumors that progress despite bevacizumab treatment. In mouse models treated with αVEGF, we demonstrated that both switching to αPD1 treatment and adding αPD1 to αVEGF treatment more effectively delayed tumor growth and prolonged survival, compared to continuing αVEGF monotherapy. Notably, αVEGF treatment is essential in preventing preserving hearing – switching to αPD1 alone and discontinuing αVEGF treatment completely abolished the hearing benefit, whereas adding αPD1 to αVEGF treatment has the most significant effect in preserving hearing. These findings suggest that αPD1 therapy may be an effective alternative treatment for patients unresponsive to bevacizumab, or those who must discontinue bevacizumab due to its side effects, while combination treatment may offer enhanced protection against tumor-induced hearing loss and tumor progression.

In summary, our study provides the first systematic characterization of the effects of ICIs on non-malignant tumors, especially the therapeutic potential of ICIs to preserve hearing. We evaluated the efficacy of αPD1 in combination with standard-of-care treatments for VS, an approach that can result in rapid translation to the clinic to improve current treatment efficacy and patient outcomes.

## Notes

### Data availability

All data and the supplementary materials from this study are included in this manuscript and are available after publication upon request from the corresponding author. Sequencing data will be deposited in GEO and will be available upon publication.

## Acknowledgments

We thank Mark Duquette and Anna Khachatryan for their superb technical support, and Dr. Peigen Huang for assisting in animal studies.

## Author contributions

L.X. designed the research and supervised the research; S. L., Z.Y., L.W., J.C. performed mouse model studies; S.L., D.C.B., L.D.L., R. S. conducted hearing test, L.M.N.W., J.L.O., and L.S.C. conducted scRNA-seq; Z.Y., Y.S., B.X., A.P.J., performed flow cytometry and histology analysis; W.H. analyzed RNASeq data; L.X., S.L., Z.Y., Y.S., A.M., analyzed data; L.X., S.R.P., L.S.C., and K.S. wrote the paper.

## Supplementary Figures

**Figure S1.**
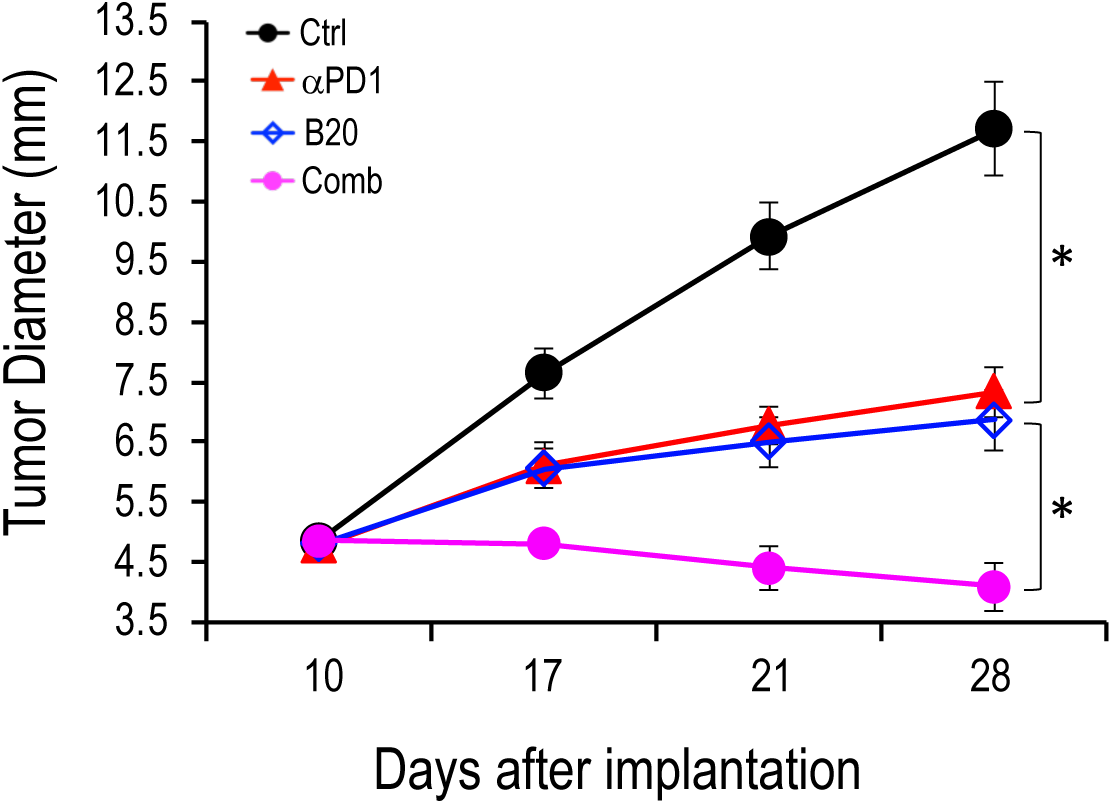
Combined αVEGF treatment enhances αPD1 efficacy in the mouse schwannoma models. Tumor diameter measured by caliper in the SC4 SN model. Animal studies are presented as mean±SEM, N=6 mice/group, and representative of at least three independent experiments. In vivo study significance was analyzed using a Student’s t-test. *P<0.05.

**Figure S2.**
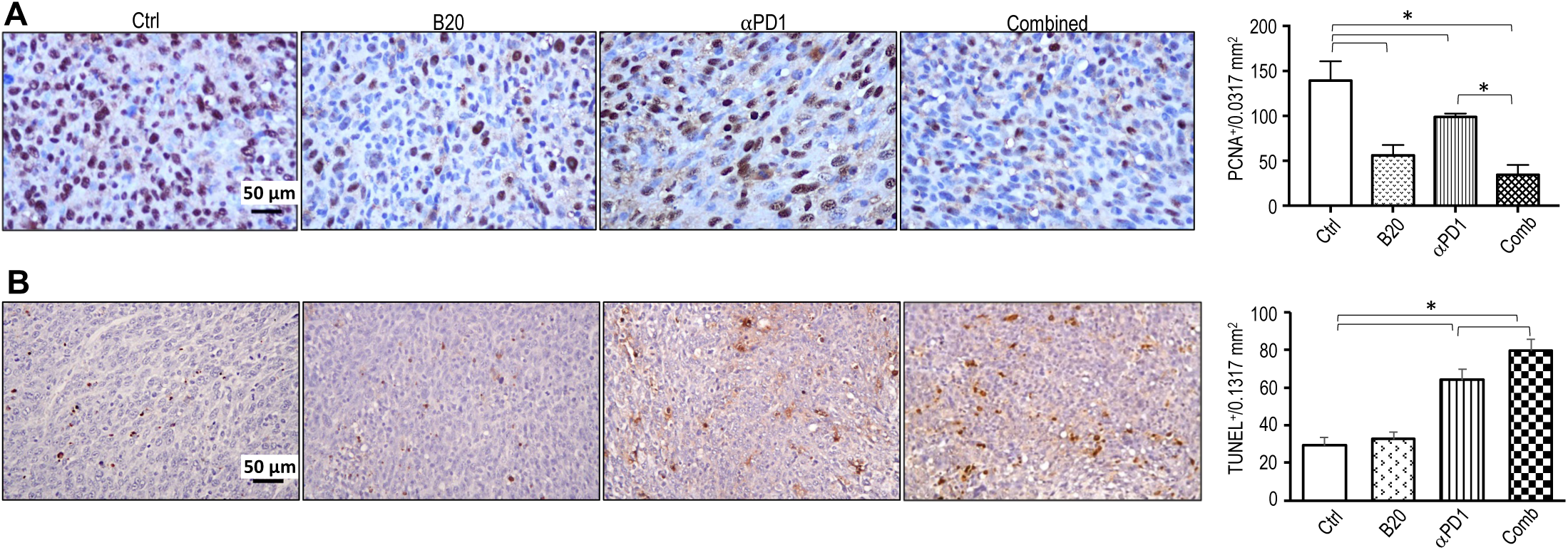
Combined αVEGF and αPD1 treatment enhance tumor control in *Nf2^-/-^*model. **(A)** Representative images of TUNEL staining for apoptotic cells **(B)** Representative images of PCNA staining for proliferating tumor cells. Image quantification and flow cytometry data are presented as mean ± SD, and analyzed using Student’s t-test and the Mann-Whitney test.

**Figure S3.**
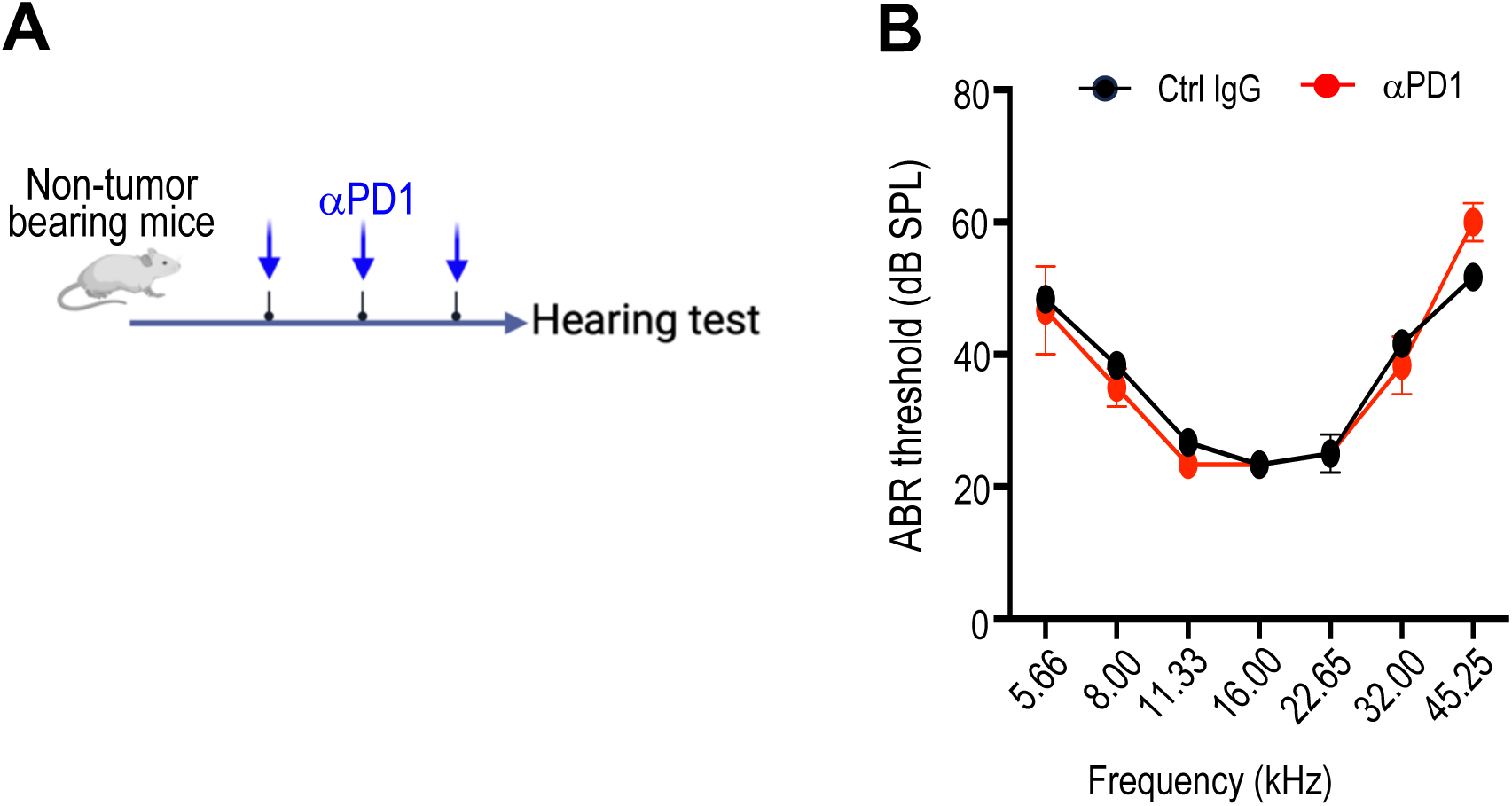
αPD1 does not cause acute ototoxicity. **(A)** Schematic of αPD1 treatment in the CPA schwannoma models. **(B)** ABR threshold of mice bearing *Nf2^-/-^* tumor in the CPA model. Animal studies are presented as mean±SEM, N=6 mice/group, and representative of at least three independent experiments. In vivo study significance was analyzed using a Student’s t-test. *P<0.05.

**Table S1.**
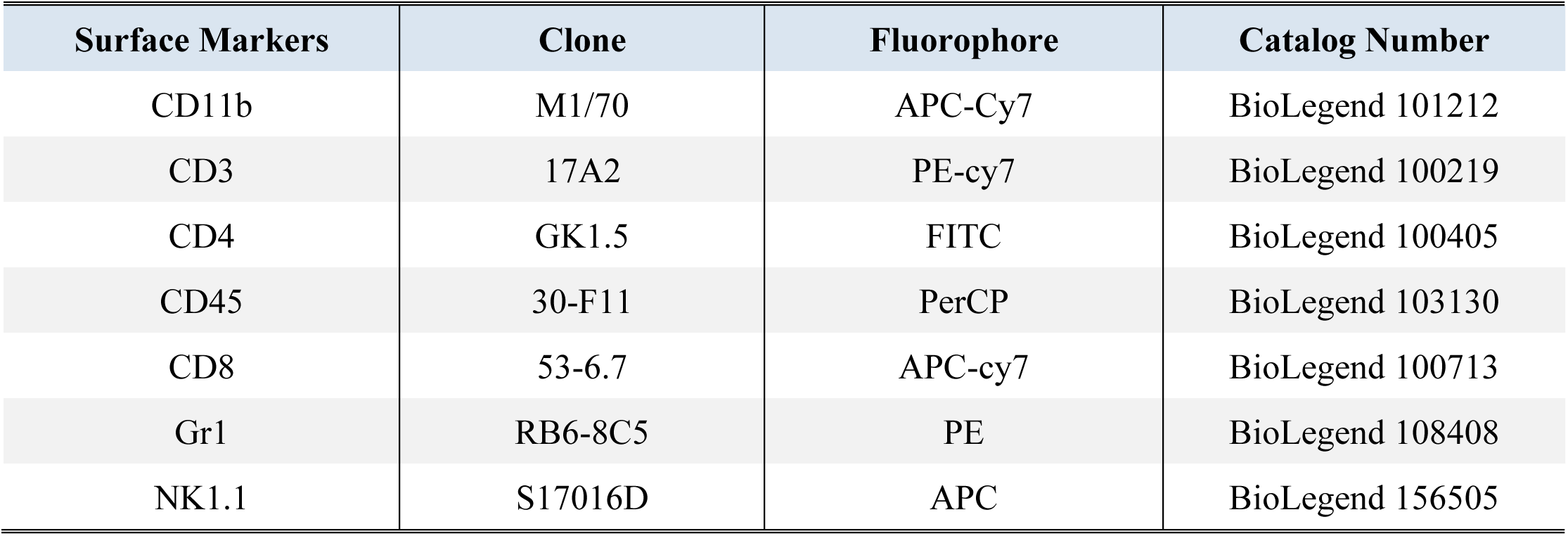
Flow cytometry antibody panels.

| Surface Markers | Clone | Fluorophore | Catalog Number |
| --- | --- | --- | --- |
| CD11b | M1/70 | APC-Cy7 | BioLegend 101212 |
| CD3 | 17A2 | PE-cy7 | BioLegend 100219 |
| CD4 | GK1.5 | FITC | BioLegend 100405 |
| CD45 | 30-F11 | PerCP | BioLegend 103130 |
| CD8 | 53-6.7 | APC-cy7 | BioLegend 100713 |
| Gr1 | RB6-8C5 | PE | BioLegend 108408 |
| NK1.1 | S17016D | APC | BioLegend 156505 |

